# The effect of microstructural variations in tendon and ligament on diffusion tensor MRI

**DOI:** 10.64898/2026.03.12.711135

**Authors:** Michael D. K. Focht, Advait Borole, Amir O. Moghaddam, Amy J. Wagoner Johnson, Roberto A. Pineda Guzman, Bruce M. Damon, Noel M. Naughton, Mariana E. Kersh

**Affiliations:** Department of Mechanical Science and Engineering, University of Illinois Urbana-Champaign, Urbana, Illinois, USA; Department of Mechanical Engineering, Virginia Tech, Blacksburg, Virginia, USA; Creare LLC, Hanover, New Hampshire, USA; Department of Biomedical and Translational Sciences, University of Illinois Urbana-Champaign, Urbana, Illinois, USA; Carl R. Woese Institute for Genomic Biology, University of Illinois Urbana-Champaign, Urbana, Illinois, USA; Carle Clinical Imaging Research Program, Stephens Family Clinical Research Institute, Carle Health, Urbana, Illinois, USA; Beckman Institute for Advanced Science and Technology, University of Illinois Urbana-Champaign, Urbana, Illinois, USA

**Author notes:** Corresponding author: *Email address:* (Mariana E. Kersh).

**Keywords:** tendon, ligament, collagen, fiber, crimp, dispersion, diffusion, DTI

## Abstract

The fibrous microstructure of tendons and ligaments is an important determinant of their mechanical behavior and integrity. Diffusion tensor imaging (DTI) is a magnetic resonance imaging (MRI) technique that enables the inference of microstructural features within fibrous tissues and has recently been used to characterize the microstructure of dense connective tissues such as tendon and ligament. However, the effect of microstructural variations in tendon and ligament on DTI metrics remains unclear. To address this gap, we simulated diffusion MRI of second harmonic generation (SHG) image-informed square lattice fiber networks to determine which microstructural features have the strongest influence on DTI metrics. Then, we performed a second set of diffusion MRI simulations for randomly dispersed fibers within synthetic tendon volumes to relate DTI metrics to the influential microstructural features, including fiber dispersion. All DTI metrics were insensitive to collagen fiber crimp. Fiber dispersion did not affect mean diffusivity, decreased axial diffusivity, increased radial diffusivity, and decreased fractional anisotropy. These results provide valuable insight into the relationships between DTI metrics and microstructural properties of tendon and ligament, which is particularly relevant for inferring microstructural changes in impaired tissue using DTI. Furthermore, our findings are an important step in the translation of DTI for clinical and computational studies of dense connective tissues such as tendon and ligament.

## 1. Introduction

Healthy tendons and ligaments are dense connective tissues primarily composed of organized collagen fibers, which work together to transmit forces within the body and support human movement [1]. As tendons and ligaments are cyclically loaded over extended periods of time, microstructural damage can accumulate in conjunction with cellular remodeling of the extracellular matrix [2]. If the accumulation of fatigue damage outpaces the remodeling of fibers, the mechanical integrity of the tissue decreases over time [3] and can lead to rupture [2, 4].

The mechanical responses of tendons and ligaments to loading are heavily influenced by their microstructural fiber organization. Blank and Roth found that increased fiber dispersion results in increased shear modulus of medial collateral ligaments, and that this effect was greater for ligaments under greater axial pretension [5]. Janssen et al. demonstrated that impaired mechanical function in the Achilles tendon resulted primarily from changes in local collagen fiber orientation and dispersion, rather than changes in fiber stiffness [6]. Thus, characterization of the heterogeneous fiber organization of tendons and ligaments is critical to accurately assess their mechanical behavior and, to be clinically feasible, must be accomplished using non-invasive methods.

Diffusion tensor imaging (DTI) is a magnetic resonance imaging (MRI) technique that enables the non-invasive assessment of water diffusion within soft tissues. Diffusion simulations have provided insight into the relationship between tissue microstructure and anisotropic water diffusion for white matter [7–9], cartilage [10–12], and skeletal muscle [13–17]. In the past 5-10 years, there has been an increase in studies using DTI to non-invasively assess tendon and ligament microstructure [18–31], including reports of correlations between DTI metrics and fiber alignment [29]. DTI metrics have also been used to detect microstructural changes due to fatigue-induced damage in both artificial fiber constructs [32] and *in vitro* tendons [31].

In addition to serving as a biomarker of tissue quality, DTI metrics can inform computational tissue models that use microstructurally-motivated constitutive parameters. For example, DTI data have been used to describe the anisotropy of fibrous tissues in computational models of tissue mechanics [33–43]. In constitutive modeling of fibrous tissues, microstructural anisotropy is often characterized by the structure tensor, which is a mathematical descriptor of the distribution of fiber orientations in threedimensional space [44, 45]. A common approach in DTI-based constitutive modeling studies has been to assume equivalence between diffusion anisotropy and tissue microstructural anisotropy [35, 36, 41]. However, water diffusion in completely aligned fiber networks also occurs perpendicular to the fibers [7, 8, 12], indicating that the diffusion tensor is more isotropic than the associated structure tensor in general.

A relationship between the anisotropy of the structure tensor and diffusion tensor for tendon and ligament has not been reported. Additionally, the anisotropy of the diffusion tensor is expected to depend on diffusion encoding parameters as well as the tissue’s MR (magnetic resonance) relaxation properties. Thus, diffusion MRI simulations incorporating realistic fiber geometries, diffusion encoding parameters, and MR relaxation properties will enable researchers to relate connective tissue microstructure to DTI metrics.

The aim of this study was to relate tendon microstructure to DTI metrics using diffusion MRI simulations with experimentally relevant diffusion encoding parameters. First, porcine digital flexor tendons were characterized using second harmonic generation (SHG) microscopy to quantify the collagen fiber microstructure. The microstructural features were varied within uniform square lattice fiber networks, and we simulated diffusion MRI to determine the sensitivity of DTI metrics to simplified tendon microstructures incorporating crimp. Based on the results of this initial sensitivity study, we performed a second set of diffusion MRI simulations for randomly dispersed fiber networks to relate the microstructural features to DTI metrics. As clinical DTI sequences continue to emerge, the relationships developed in this study will aid in i) the interpretation of DTI metrics for clinical assessments and ii) the implementation of DTI-based computational models of tendon and ligament function.

## 2. Methods

### 2.1 Tendon samples, SHG imaging, and microstructural measurements

Porcine digital flexor tendons (n=9) were harvested, and SHG images were collected in the mid-substance of the tendon (Figure 1A). The SHG images were collected as part of another study [31] and include a mixture of fatigued and non-fatigued tendons. The cryosectioned tendons were imaged (1.66 *µ*m pixel resolution) using a Zeiss LSM 710 multiphoton microscope (Zeiss, Oberkochen, Germany) equipped with a MaiTai DeepSee Ti:Sapphire femtosecond laser (Spectra-Physics, Newport Crop., USA). Ten randomly sampled 500×2000 *µ*m regions of interest (ROI) were identified and collagen fiber diameter, spacing between fibers, crimp wavelength, and crimp height were measured at five random locations within each ROI, similar to our previous studies [31, 46, 47]. Fiber area fraction was computed by binarizing (ImageJ Software; NIH, Bethesda, MD, USA) each 500×2000 *µ*m image region (n=10) and dividing the number of pixels assigned to the fibers by the total number of pixels.

**Figure 1:**
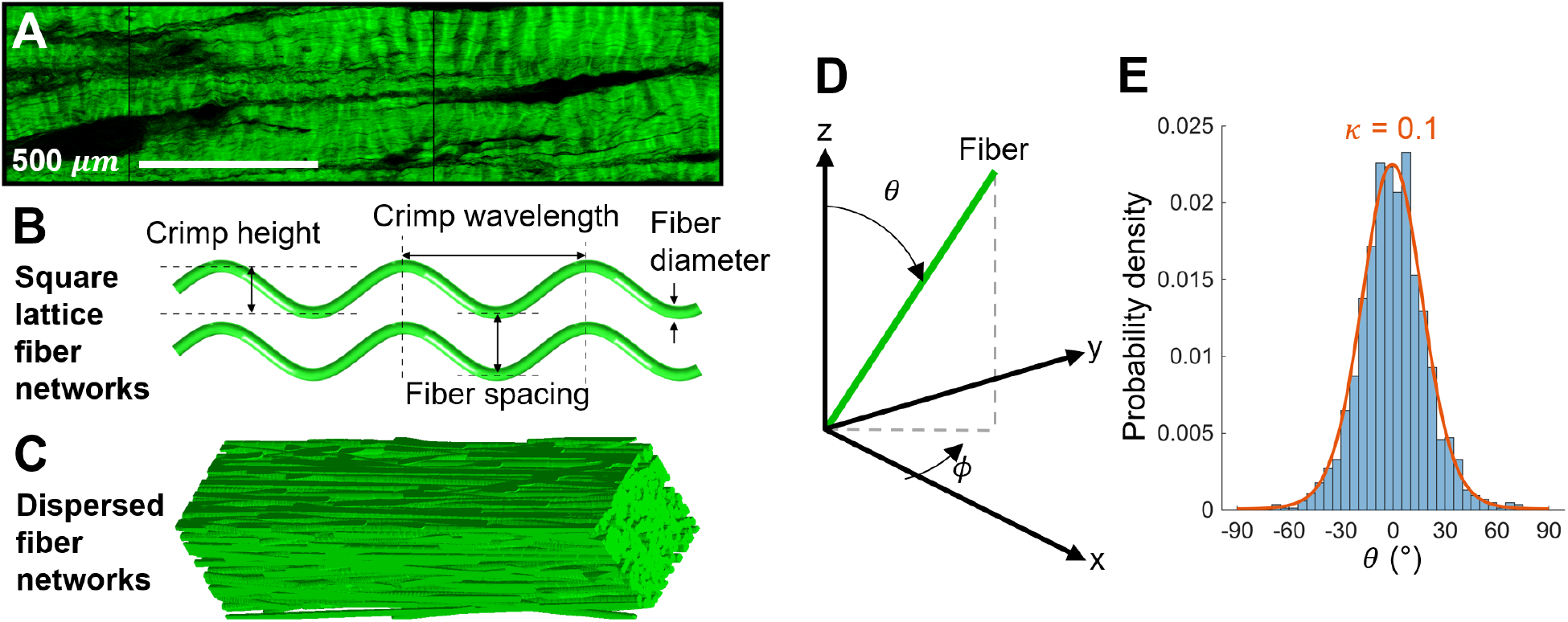
Microstructurally-informed models of tendon for diffusion MRI simulations. A) Representative second harmonic generation image of tendon. B) Microstructural parametrization of the square lattice fiber networks. C) Representative dispersed fiber network. D) Coordinate system used to define the fiber orientation with respect to the longitudinal (z) axis and transverse (x–y) plane in the dispersed fiber networks. E) Representative histogram of *θ*, the angle that a fiber makes with the z axis in the dispersed fiber networks, with the analytical von Mises distribution (orange curve) overlaid for *κ* = 0.1.

### 2.2 Diffusion MRI simulations

Diffusion MRI simulations were performed over microstructural tendon domains of collagen fibers using a previously reported hybrid lattice Boltzmann method [48] (Python) to simulate the Bloch-Torrey equation. For all simulations, a pulsed gradient spin echo (PGSE) pulse sequence was simulated with a diffusion gradient duration (*δ*) of 4 ms, diffusion gradient separation (Δ) of 10 ms, and b-value of 400 s/mm^2^. An image formation time of 4 ms was added to the end of the sequence yielding a simulation echo time (TE) of 18 ms, at which point the signal within the volume was integrated to yield a voxel-averaged signal. 12 uniformly spaced gradient directions were simulated for each microstructural geometry along with one non-diffusion weighted sequence and the resulting diffusion-weighted signal intensities and non-diffusion-weighted signal were used to compute the diffusion tensor and its associated eigenvalues (*λ*_1_, *λ*_2_, *λ*_3_). Four DTI metrics were then calculated from the diffusion tensor: mean diffusivity (MD = (*λ*_1_ + *λ*_2_ + *λ*_3_)*/*3), axial diffusivity (AD = *λ*_1_), radial diffusivity (RD = (*λ*_2_ + *λ*_3_)*/*2), and fractional anisotropy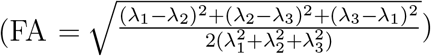.

Tendon has four discrete proton populations that act as sources of MR signal (Figure 2). However, two of these populations, tropocollagen and bound water, exhibit very low T_2_ values that cause their associated signal to rapidly decay with minimal contribution to the final MR signal. As such, we only explicitly considered the signal contribution of the interfibrillar water within fibers and interstitial water between fibers in our simulations (Table 1). The interfacial boundaries between collagen fibers and interstitial water were modeled as impermeable membranes ([49]).

**Table 1:** MR relaxation parameters used for all simulations. Each fiber network domain comprised two subdomains: collagen fibers and interstitial water.

|  | Collagen fibers | Interstitial water |
| --- | --- | --- |
| Diffusivity ($\mu\text{m}^2/\text{ms}$) | 1 [51] | 2 [55] |
| $T_2$ (ms) | 20 [49, 50] | 62 [49, 50] |
| Water fraction | 0.22 [49, 50] | 1 |

**Figure 2:**
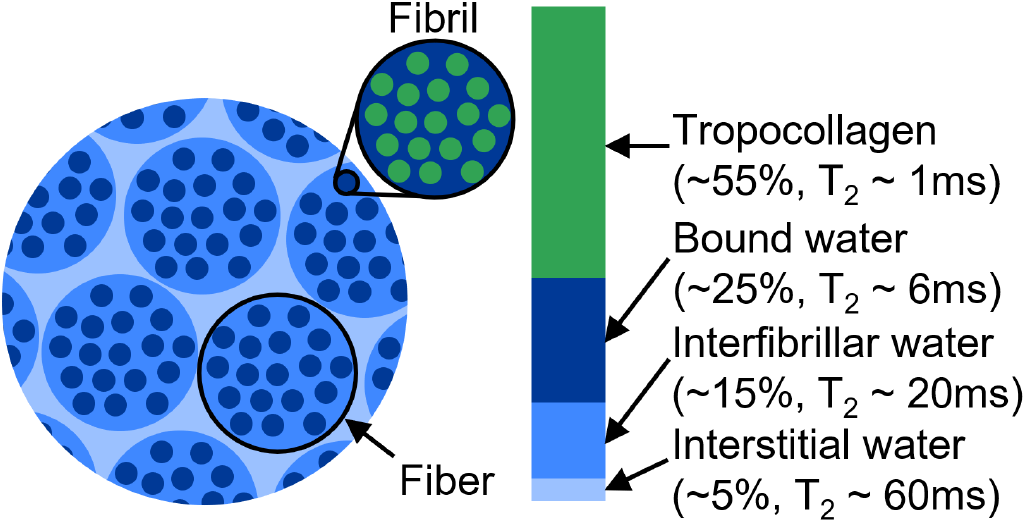
Tendon proton populations contributing to MR signal [49–51]. Percentages indicate the fraction of the total MR signal attributable to each proton population. The fibers in the diffusion MRI simulations, as labeled here, are defined as bundles of collagen fibrils.

Two classes of microstructural domains were examined. The first consisted of simplified biphasic square lattice domains to identify the sensitivity of DTI signal to changes in microstructural features, while the second consisted of voxel-scale domains that accounted for fiber orientation and dispersion.

All simulations were performed using Virginia Tech’s Advanced Research Computing’s Tinker Cliffs and Owl computing clusters.

### 2.3 Square lattice fiber network sensitivity study

To determine how changes in tendon microstructure impact DTI measurements a global sensitivity analysis was performed [52]. We adopt the method developed by Saltelli et al. [53, 54], which quantifies how much of the variance of a model’s output can be attributed to variance of individual input parameters or non-linear combinations of multiple parameters. First-order sensitivity indices describe the fraction of the total variance of the output attributed to the variance of a particular parameter. For example, a firstorder fiber spacing sensitivity index of 0.75 for RD would mean that 75% of the total variance in RD is attributable to variance in the fiber spacing. Second-order indices capture the response of the model to the combination of two parameters that cannot be written as a superposition of two first-order effects independently. Total-order indices are the sum of first, second, and higherorder indices for each parameter and describe the total contribution of a parameter to the variance of the system output.

Three-dimensional biphasic domains were generated comprising of crimped collagen fibers aligned in a square lattice and embedded in interstitial water (Figure 1B). Crimp of the collagen fibers was modeled as a sine wave. Domains were parameterized by collagen fiber diameter, fiber spacing, crimp wavelength, and crimp height. Parameter ranges were defined over physiologically realistic bounds based on the microstructural features measured in the SHG images and reported in Table 2. Based on the parameterized model, a computationally efficient unit cell consisting of a single collagen fiber over one crimp wavelength with periodic boundary conditions, appropriately modified to account for phase shift in the MR signal [48, 56], was defined.

**Table 2:** Second harmonic generation-based measurements of the microstructural parameters and the parameter ranges used for the sensitivity study of the square lattice fiber networks. Fiber area fraction was only used to inform the volume fraction range of the dispersed fiber networks and was not used to parameterize the square lattice fiber networks for the sensitivity study.

| | Fiber<br>diameter<br>($\mu\text{m}$) | Fiber<br>spacing<br>($\mu\text{m}$) | Crimp<br>wavelength<br>($\mu\text{m}$) | Crimp<br>height<br>($\mu\text{m}$) | Fiber<br>area<br>fraction |
| --- | --- | --- | --- | --- | --- |
| Mean | 51 | 10.6 | 64.3 | 10.9 | 0.872 |
| Standard deviation | 22.3 | 4.3 | 15.9 | 3.2 | 0.069 |
| Sensitivity study range | 20–80 | 5–15 | 50–80 | 5–15 | N/A |

A Sobol sampling method [53, 57], which provides low-discrepancy, quasi-random sampling, was then used to sample the microstructural parameter space. Parameters were sampled independently from a uniform distribution over each parameter range. Sobol sampling yielded 2N(p+1) parameter combinations, where p=4 is the number of parameters and N=1024 is the number of independent samples of each parameter. This resulted in the generation of 10,240 microstructurally unique unit cell domains over which DTI simulations using the sequence described above were performed. Because unit cell domain sizes varied based on fiber diameter and spacing, unit cells were dynamically spatially discretized such that between 40 and 60 nodes spanned each cross-section direction of the unit cell, yielding dx values of ∼1 *µ*m. Temporal discretization was then determined based on the selected dx using a previous heuristic scaling that was found to minimize numerical error [15], yielding dt values of 20–125 *µ*s. Simulation results were then used to compute global sensitivity indices using the open source SaLib python package [58].

### 2.4 Dispersed fiber networks

The second set of domains incorporated fiber dispersion within synthetic MRI voxels of tendon (Figure 1C). Domains consisted of collagen fibers embedded in interstitial water and oriented to match an overall dispersive organization as characterized by the dispersion parameter *κ*, which we adopt from the Holzapfel-Gasser-Ogden (HGO) model for fibrous materials’ mechanical behavior [45]. In the HGO model, the orientation of fibers is transversely isotropic and *κ* ranges from 0 (fully anisotropic) to 1/3 (fully isotropic). To generate domains, fibers were placed in the domain with a random seed point and angle in the transverse (x–y) plane, *ϕ*, sampled from a uniform distribution (Figure 1D). The angle between the fiber and the longitudinal (z) axis, *θ*, was prescribed by randomly sampling from a von Mises distribution corresponding to the dispersion parameter, *κ* (Figure 1E). Fibers were modeled as straight cylinders (due to DTI’s low sensitivity to crimp in the square lattice fiber networks; see Figure 3) and allowed to intersect.

**Figure 3:**
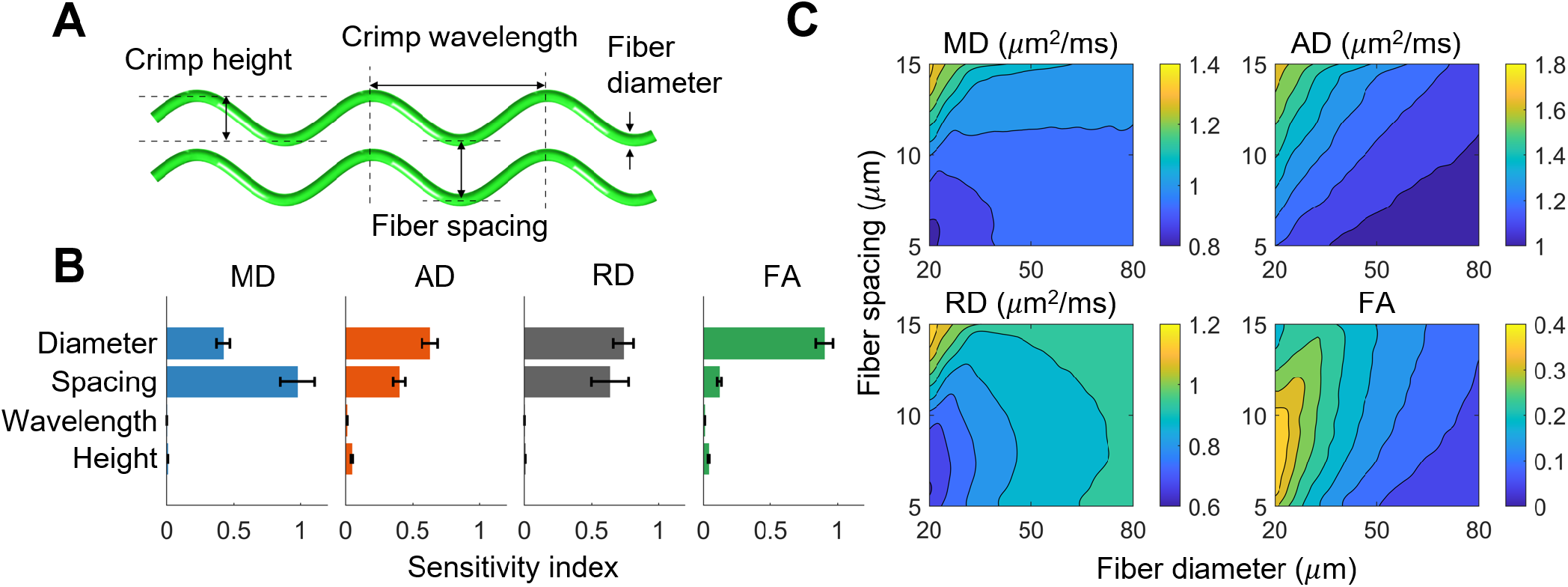
Sensitivity study results for the square lattice fiber networks. A) Microstructural parametrization of the square lattice fiber networks. B) Sobol total sensitivity indices. Error bars indicate 95% confidence intervals. C) Contour plots of DTI metrics vs fiber diameter and spacing. Crimp parameters were averaged out, because all DTI metrics were insensitive to crimp.

Domains were defined for five levels of fiber diameter (30–70 *µ*m), five levels of volume fraction (0.75–0.95), and ten levels of dispersion, *κ* (0– 1/3), leading to 250 different microstructural parameter combinations. For each combination, five separate domains were generated to account for the random nature of the fiber geometries (MATLAB R2024a; Mathworks, Natick, MA, USA). Domains were 500×500×2000 *µ*m to approximate voxel dimensions achievable on preclinical MRI scanners and had a spatial resolution of 1 *µ*m and temporal resolution of 111.1 *µ*s. Phase-shift adjusted periodic boundary conditions were applied on all sides. DTI simulations were then performed over the domains using the sequence described above, DTI metrics were computed, and the mean and standard deviation of each DTI metric was calculated for each microstructural parameter combination.

### 2.5 Modeling effect of fiber dispersion on FA

Prior investigations have considered the impact of fiber dispersion on DTI measurements in fibrous tissues such as cartilage and neural white matter [35, 36, 41]. These studies have proposed a mapping that connects the diffusion tensor to the dispersion parameter *κ* such that fully isotropic fiber arrangements (*κ*=1/3) correspond to isotropic diffusion behavior (AD=RD; which assumes transversely isotropic diffusion) while perfectly anisotropic fiber arrangements (*κ*=0; e.g. fibers completely aligned in axial direction) correspond to completely anisotropic diffusion (AD ≫RD; which implies RD→0). This yields a relationship between the axial and radial diffusivity of AD/RD = (1−2*κ*)*/κ*. Incorporating this relationship into the definition of FA then yields FA = (*β*−1)*/*(*β*^2^+2)^1*/*2^ where *β* = AD/RD. Directly mapping fiber dispersion to diffusion anisotropy notably implies that the radial diffusivity becomes negligible for highly aligned fibers. Such a result may be plausible in the long diffusion time limit of fibrous tissues with small fibers such as white matter. However, in tendon, the fibers are much larger and diffusion times must be kept relatively short to preserve signal, which provides a pathway for radial diffusion even when fibers are perfectly aligned in the axial direction (*κ*=0).

To incorporate this radial diffusion behavior in a simple analytical model, we define a new mapping between fiber dispersion and diffusion anisotropy that is similar to the direct mapping model. However, for fully anisotropic fiber arrangements, radial diffusion is modeled using the Maxwell-Garnett, which provides a first-order approximation of diffusion in insulated cylinders [59, 60]. We refer to this model as the dispersed Maxwell-Garnett. This yields a new relationship between the axial and radial diffusivity of AD/RD = 3*κ* + (1 + *v*)(1− 3*κ*) where *v* is the collagen volume fraction. This then yields FA = (*α* − 1)*/*(*α*^2^ + 2)^1*/*2^ where *α* = AD/RD.

## 3. Results

### 3.1 SHG microscopy

Collagen fibers had a mean diameter of 51 *µ*m, mean spacing of 10.6 *µ*m, mean crimp wavelength of 64.3 *µ*m, and mean crimp height of 10.9 *µ*m (Table 2), as measured experimentally by SHG. The distributions of these microstructural features were used to inform the microstructural parameter ranges for the square lattice fiber network simulations. Collagen fiber area fraction had a mean value of 0.872. The distributions of fiber diameter and area fraction were used to inform the range of fiber diameters and volume fractions simulated in the dispersed fiber networks.

### 3.2 Square lattice fiber network sensitivity study

DTI metrics were orders of magnitude (10– 500x) less sensitive to crimp height and wavelength (range of Sobol total sensitivity values: 0.001–0.047) than fiber diameter and spacing (range: 0.118–0.977) (Figure 3B). For example, MD was approximately twice as sensitive to fiber spacing as it was to diameter (0.977 vs 0.421). In contrast, AD, RD, and FA were more sensitive to fiber diameter (0.629, 0.739, and 0.901, respectively) than spacing (0.398, 0.637, and 0.118).

Interestingly, AD is over seven times more sensitive to crimp height than RD (0.047 vs 0.006). All DTI metrics were highly insensitive to crimp wavelength, with a maximum sensitivity of 0.011 for AD.

There are several interactions between parameters worth noting, as quantified by second-order Sobol sensitivity indices. All DTI metrics were most sensitive to fiber spacing for small fiber diameters (see increased density of contours for small diameters in Figure 3C). MD, AD, and RD were most sensitive to interactions between fiber diameter and spacing (second-order sensitivities of 0.39, 0.068, and 0.377, respectively). AD increased as diameter decreased and spacing increased in a more consistent manner than the other metrics (see linear contour lines in Figure 3C, top right). RD is the driver of the highly nonlinear behavior present in MD and FA, as both depend on RD. Finally, FA decreased as diameter increased, with a local maximum (0.39) occurring at a spacing of ∼7.5 *µ*m for small diameters (20 *µ*m). FA was most sensitive to diameter for small spacings (second-order sensitivity of 0.067). Neither crimp wavelength nor height had strong interactions with other microstructural parameters. RD had the highest crimp-related second-order sensitivity with a value of 0.019 between crimp height and fiber spacing.

### 3.3 Dispersed fiber networks

As the fiber networks became increasingly dispersed (Figure 4A), MD did not change, AD decreased, RD increased, and FA decreased (Figure 4B). MD increased as fiber diameter increased and volume fraction decreased and was unaffected by dispersion. Interestingly, AD did not depend on fiber diameter for highly aligned fiber networks and increasingly depended on fiber diameter as dispersion increased. This effect is likely due to fiber boundaries only restricting radial diffusion when there is no dispersion. However, as fibers become increasingly dispersed, they also start restricting diffusion in the axial direction. As they do so, smaller fiber diameters result in more fiber boundaries for a given volume fraction, thus resulting in more restricted axial diffusion for a given volume fraction and dispersion level. RD depended on all microstructural features and increased as fiber diameter increased, volume fraction decreased, and dispersion increased. For a given microstructural parameter combination, the standard deviation was lowest for AD, which implies that it was the least sensitive DTI metric to random variations in fiber network geometry.

**Figure 4:**
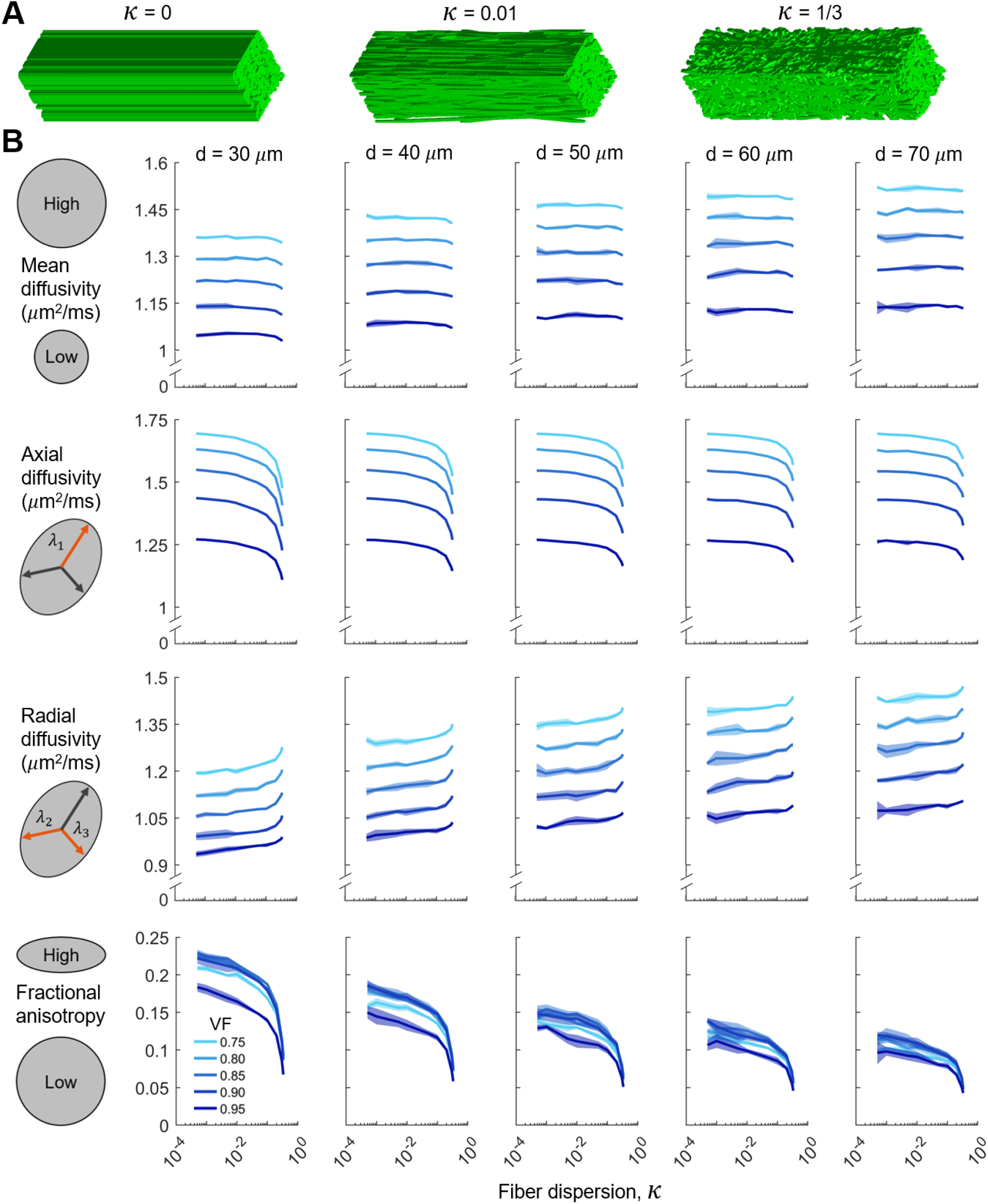
Simulation results for the dispersed fiber networks. A) Representative dispersed fiber networks with increasing fiber dispersion, *κ*. B) Relationships between the DTI metrics and the fiber dispersion, *κ*, for the dispersed fiber networks with varying fiber diameter (d) and volume fraction (VF). The shaded regions around the mean curve indicate one standard deviation from the mean in the positive and negative directions. Semi-log axes are used to emphasize the behavior for low levels of dispersion.

Similar to the square lattice simulations, the relationship between FA and volume fraction was non-monotonic, with a local maximum occurring between a volume fraction of 0.85 and 0.9 (Figure 5). This local maximum is likely caused by the reduction in fiber boundaries as volume fraction increases beyond 0.85–0.9. In such a case, the domain approaches a contiguous block of collagen, which, in our simulations, has increasingly isotropic diffusion properties as the amount of fiber boundaries decrease. FA decreased as fiber diameter increased because there are fewer fiber boundaries to restrict radial diffusion as fiber diameter increases. Finally, FA decreased as dispersion increased (Figures 4 and 5).

**Figure 5:**
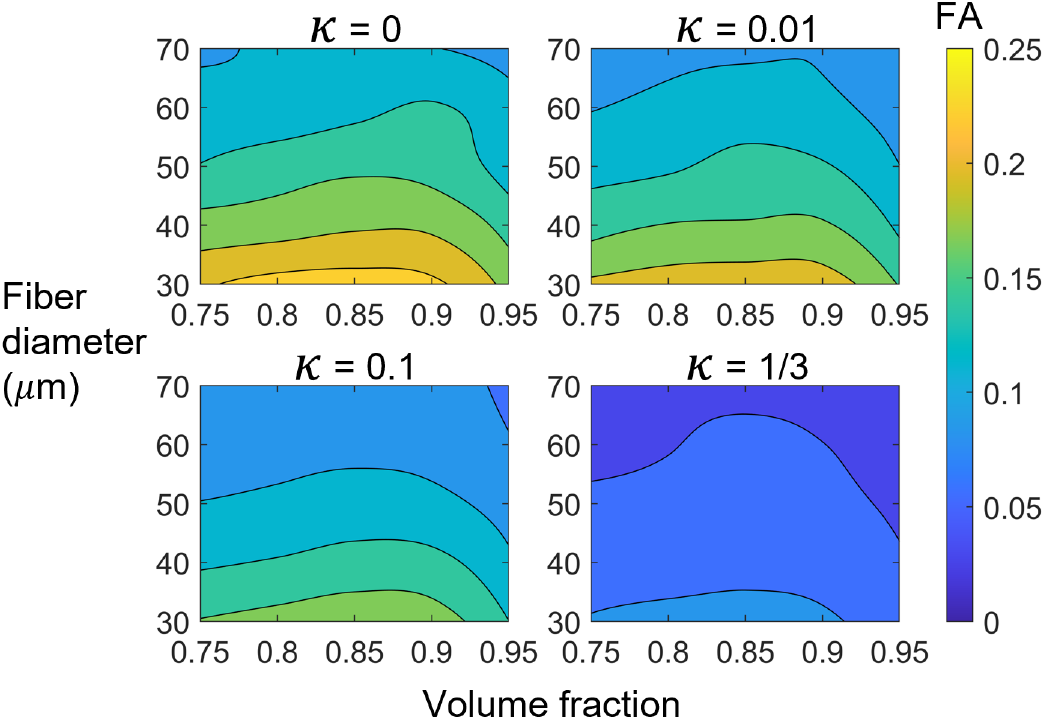
Contour plots of fractional anisotropy (FA) vs fiber diameter and volume fraction for four levels of dispersion, *κ*.

### 3.4 Effect of fiber dispersion on FA

The simulated FA-*κ* relationship did not match the direct mapping model or dispersed Maxwell-Garnett model (Figure 6). When *κ*=0, the simulated FA was ∼0.15 in contrast to the direct mapping value of 1 and Maxwell-Garnett value of ∼0.35. As *κ* approached 1/3 (i.e., increasingly isotropic microstructure), the difference between the simulated, direct mapping, and dispersed Maxwell-Garnett FA values decreased. Unlike the direct mapping and dispersed Maxwell-Garnett models, our simulations did not achieve complete diffusion isotropy (FA=0). This is likely due to boundary effects, since the domains become increasingly aperiodic as the fibers become more dispersed.

**Figure 6:**
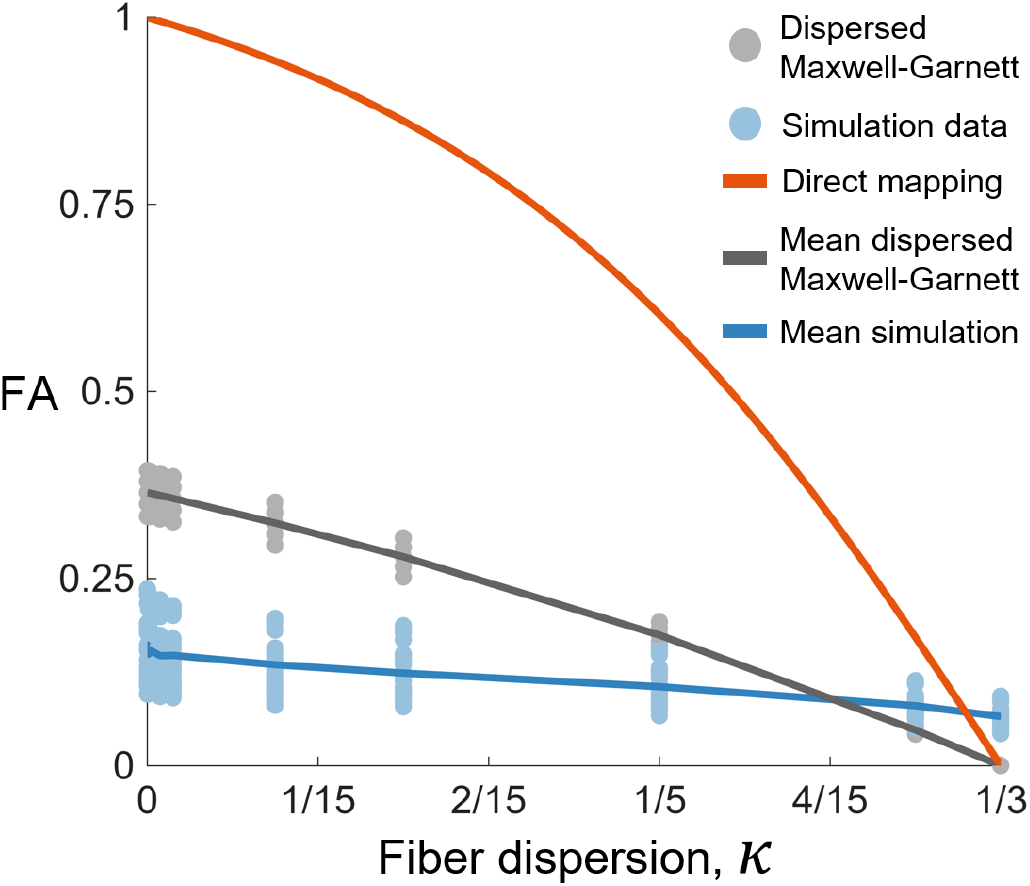
The relationship between fractional anisotropy (FA) and fiber dispersion, *κ*, from the direct mapping model (orange), dispersed Maxwell-Garnett model (gray), and the simulations (blue). The mean dispersed Maxwell-Garnett curve is the result for a volume fraction of 0.85 and the mean simulation curve is the result for a volume fraction of 0.85 and fiber diameter of 50 *µ*m.

## 4. Discussion

In order to improve the interpretability of DTI for dense connective tissues such as tendon and ligament, we simulated diffusion MRI of synthetic collagen fiber networks and quantified the relationship between the tissue microstructure and DTI metrics. Our finding that DTI metrics were insensitive to fiber crimp (Figure 3) implies that crimp morphologies are unable to be effectively inferred using DTI. The simulated diffusion tensors were substantially more isotropic than the fiber microstructures (Figure 6), and the relationship between fractional anisotropy and fiber dispersion depended on both fiber diameter and volume fraction (bottom of Figure 4).

Our results highlight the importance of using tissue-specific relationships between fractional anisotropy and fiber dispersion (Figure 6). The simulated FA for tendon did not follow the direct mapping or dispersed Maxwell-Garnett curves. According to our diffusion MRI simulations of tendon, the structure tensor, which describes the microstructural orientation and dispersion of fibers, is substantially more anisotropic than the diffusion tensor (blue curve ≪orange curve in Figure 6). Thus, the direct mapping and dispersed Maxwell-Garnett models result in an underestimation of the microstructural anisotropy of tendon for a given FA, although the dispersed Maxwell-Garnett model is a marked improvement compared to the direct mapping model. The dispersed Maxwell-Garnett model assumes long diffusion times and therefore neglects any signal from the water inside the fibers (intrafibrillar water in this case). This assumption is valid for white matter, which has axon fiber diameters of 1 ∼*µ*m. However, tendon collagen fibers are ∼50 *µ*m in diameter, and diffusion times must be kept relatively short in order to preserve signal. Thus, for realistic diffusion times, the long diffusion time assumption does not hold for tendon. This is further illustrated by the fact that the dispersed Maxwell-Garnett model does not depend on fiber diameter, whereas the FA-*κ* relationship from the simulations does depend on fiber diameter (see bottom of Figure 4)

Our dispersed fiber network simulations resulted in slightly lower diffusivity and higher FA when compared to experiments using the same diffusion encoding parameters [18, 31]. For example, for a volume fraction of 0.85, fiber diameter of 50 *µ*m, and *κ* of 0.05, the simulated MD and FA were 1.31 *µ*m^2^/ms and 0.13 in comparison to 1.53 *µ*m^2^/ms and 0.08 from experimental measurements. This reasonably close agreement between our simulation results and experiment is despite our choice of diffusion coefficients from previous literature. Additionally, the lower FA in the experimental data may be influenced by the regional voxel averaging used in the experimental study, which tends to lower FA by averaging together signals from voxels with differing principal fiber orientations.

The square lattice fiber network simulations of the mean microstructure (50 *µ*m diameter and 10 *µ*m spacing) resulted in lower diffusivity (MD=0.98 *µ*m^2^/ms) and higher FA (0.16) compared to that of the dispersed fiber networks mentioned previously. This discrepancy is primarily due to the dispersed fiber networks having intersecting fibers and more dense fiber packing than the square lattice fiber networks. Although the square lattice arrangement of fibers enables an efficient analysis of model sensitivities to simple microstructural features such as crimp, it does not allow for physiologically realistic volume fractions or fiber dispersion. For example, the square lattice fiber arrangement has a maximum possible volume fraction of ∼0.785 [59], whereas the mean area fraction of the SHG images was 0.872 (Table 2). Furthermore, the intersecting fibers of the dispersed fiber networks may be more physiologically realistic than the non-intersecting fibers of the square lattice fiber networks, as fibers were observed to split and merge in the SHG images.

In agreement with our results, previous studies utilizing fiber phantoms and water diffusion simulations found that diffusion anisotropy is higher for more aligned fiber networks [10, 11, 61]. However, others have reported that FA is directly proportional to fiber density [7–9], which is not in complete agreement with our results. We found that FA increased with volume fraction fairly linearly until reaching a critical value between 0.85 and 0.9, after which FA decreased. Notably, this critical value is near the mean area fraction of 0.872 measured in the SHG images (Table 2). The non-monotonic relationship between FA and volume fraction is due to our simulations’ use of high volume fractions and intersecting fibers, combined with the inclusion of interfibrillar (within-fiber) water diffusion. As volume fraction increased beyond 0.85–0.9, there were large contiguous regions of collagen and thus fewer fiber boundaries to restrict radial diffusion. This effect is likely a limitation of the model, as ∼0.907 [59] is the limit of volume fraction for non-intersecting packed cylinders of uniform diameter. Notably, excluding the results for a volume fraction of 0.95, there is relatively small effect of volume fraction on FA as compared with diameter (see shallow slope of contour lines to the left of volume fraction = 0.9 in Figure 5).

Our findings that i) diffusivity decreased as volume fraction increased and ii) FA decreased as fiber dispersion increased (Figure 4B) conflict with Achilles tendon 3T data [29] wherein diffusivity and FA were positively correlated with collagen content and fiber dispersion, respectively. This surprising result, also acknowledged by the authors, may be a result of the specific imaging protocol used. In their work, the tendon was positioned at an angle of 55 degrees with respect to B_0_ to utilize the magic angle effect [62, 63], and they used a TE of 36 ms. The magic angle effect greatly increases the T_2_ of the water in close contact with the tropocollagen molecules, and therefore results in much higher signal from the bound water (diffusivity of ∼0.6 *µ*m^2^/ms, [64]). This effect may explain the lower diffusivity measurements reported in their study (MD=0.71 *µ*m^2^/ms). Furthermore, the authors hypothesized that the polarized light method used for quantifying microstructural anisotropy may have been assessing fiber dispersion at a different length scale than diffusion MRI.

These relationships between DTI metrics and tendon microstructure are particularly valuable for inferring microstructural changes in impaired tissue. Tendon disuse leads to more dispersed collagen fibers [6], and chronic tendinopathy has been associated with localized regions of decreased collagen density and increased fiber dispersion [65]. Similarly, acutely injured tendon has more dispersed collagen fibers than uninjured tendon [66]. Interestingly, both increased collagen fiber dispersion and decreased collagen volume fraction increase RD and decrease FA in our simulations. More work is needed to experimentally confirm these predictions.

The effect of noise was not considered in this study and is the subject of ongoing work. Clinical studies of tendon and ligament that use DTI face the challenge of low signal-to-noise ratio, which is important to consider when interpreting experimental results. Despite this challenge, DTI is a promising method for non-invasively assessing the microstructure of tendon and ligament, and our results provide insight into how DTI metrics are expected to change as a result of microstructural changes. Non-invasive methods for assessing tendon and ligament microstructure, such as DTI, are needed in order to improve clinical outcomes and inform computational models of tissue mechanics.

## 5. Conclusion

This work provides microstructural interpretations for DTI of tendon and ligament and highlights the utility of simulating diffusion MRI when using DTI to infer microstructural properties of tissue. DTI metrics are insensitive to collagen fiber crimp; thus, crimp properties are unable to be effectively inferred with the diffusion encoding parameters simulated in this study. Generally, greater collagen fiber volume fraction decreases apparent diffusivity. Collagen fiber dispersion does not affect MD, decreases AD, increases RD, and decreases FA. Microstructural anisotropy is not equivalent to diffusion anisotropy in tendon and ligament for clinically relevant diffusion encoding parameters, as the diffusion tensor is substantially more isotropic than the underlying microstructural fiber distribution. With the aid of tissue- and scan-specific simulations, DTI can be used to non-invasively estimate microstructural properties of tissue and inform constitutive models for finite element simulations of tissue mechanics. Overall, these results provide valuable insight into the relationship between DTI metrics and the microstructure of tendon and ligament.

## 6. Acknowledgments

The authors thank Dr. Shreyan Majumdar for helping to develop the pulse sequence parameters used in this study.

The authors acknowledge support from NIH/NIAMS 1 R01 AR073831 and the Stephens Family Clinical Research Institute.

## 7. Conflict of Interest

Declarations of interest: none.

## Notes

https://doi.org/10.13012/B2IDB-9327074_V1

### Competing Interest Statement

The authors have declared no competing interest.

